# Tumor Subtype Classification Tool for HPV-associated Head and Neck Cancers

**DOI:** 10.1101/2024.07.05.601906

**Authors:** Shiting Li, Bailey F. Garb, Tingting Qin, Sarah Soppe, Elizabeth Lopez, Snehal Patil, Nisha J. D’Silva, Laura S. Rozek, Maureen A. Sartor

**Author notes:** Corresponding Authors: Maureen A. Sartor, University of Michigan, 2017 Palmer Commons, Ann Arbor, MI 48109-2218.; and Laura S. Rozek, Georgetown University, 2115 Wisconsin Ave NW, Washington DC 20009. Co-first authors (Shiting Li, Bailey F. Garb). Georgetown University, Oncology Department, School of Medicine, Washington DC, USA.

## Abstract

**Importance:** Molecular subtypes of HPV-associated Head and Neck Squamous Cell Carcinoma (HNSCC), named IMU (immune strong) and KRT (highly keratinized), are well-recognized and have been shown to have distinct mechanisms of carcinogenesis, clinical outcomes, and potentially differing optimal treatment strategies. Currently, no standardized method exists to subtype a new HPV+ HNSCC tumor. Our paper introduces a machine learning-based classifier and webtool to reliably subtype HPV+ HNSCC tumors using the IMU/KRT paradigm and highlights the importance of subtype in HPV+ HNSCC.

**Objective:** To develop a robust, accurate machine learning-based classification tool that standardizes the process of subtyping HPV+ HNSCC, and to investigate the clinical, demographic, and molecular features associated with subtype in a meta-analysis of four patient cohorts.

**Data Sources:** We conducted RNA-seq on 67 HNSCC FFPE blocks from University of Michigan hospital. Combining this with three publicly available datasets, we utilized a total of 229 HPV+ HNSCC RNA-seq samples. All participants were HPV+ according to RNA expression. An ensemble machine learning approach with five algorithms and three different input training gene sets were developed, with final subtype determined by majority vote. Several additional steps were taken to ensure rigor and reproducibility throughout.

**Study Selection:** The classifier was trained and tested using 84 subtype-labeled HPV+ RNA-seq samples from two cohorts: University of Michigan (UM; n=18) and TCGA-HNC (n=66). The classifier robustness was validated with two independent cohorts: 83 samples from the HPV Virome Consortium and 62 additional samples from UM. We revealed 24 of 39 tested clinicodemographic and molecular variables significantly associated with subtype.

**Results:** The classifier achieved 100% accuracy in the test set. Validation on two additional cohorts demonstrated successful separation by known features of the subtypes. Investigating the relationship between subtype and 39 molecular and clinicodemographic variables revealed IMU is associated with epithelial-mesenchymal transition (p=2.25×10^−04^), various immune cell types, and lower radiation resistance (p=0.0050), while KRT is more highly keratinized (p=2.53×10^−08^), and more likely female than IMU (p=0.0082).

**Conclusions and Relevance:** This study provides a reliable classifier for subtyping HPV+ HNSCC tumors as either IMU or KRT based on bulk RNA-seq data, and additionally, improves our understanding of the HPV+ HNSCC subtypes.

## Introduction

Cancer types are typically categorized according to the cell of origin, but vast heterogeneity often exists within these groupings^1^. These groupings often have clinical utility as prognostic biomarkers, aid physicians in therapeutic strategies, and are frequently associated with treatment response ^2^. Research in breast ^2^, lung ^3^, pancreatic ^4^ and colon ^5^ cancers has uncovered distinct subtypes based on specific gene driver mutations or epigenetic signatures. By integrating these and additional diverse data sources, clear pathways of tumor development can be established ^6^. As an alternative to this attribute-based approach, bulk RNA-seq has been widely used to define cancer subtypes by clustering ^7^. As precision medicine and targeted therapies advance, the utility of defining more narrowly-defined subtypes is amplified.

HPV-associated head and neck cancer continues to increase at an epidemic level. Approximately 30% of HNSCC ^8^ can be attributed to human papillomavirus (HPV) with oropharyngeal (OPSCC) being the most common site associated with HPV ^9^. HPV infection currently drives approximately 71% and 52% of OPSCC in the USA and UK, respectively, ^10^ and typically confers a survival advantage, with 5-year survival rates averaging ∼80% ^11^. While there is a wide morphologic and epigenetic ^12^ diversity within HPV+ HNSCC, tumor subtyping is not yet widely used for this cancer population.

HPV+ HNSCC molecular subtyping has been conducted by multiple groups as reviewed by Qin et al^11^. Most of these studies used gene expression levels to define the subtypes ^13–15^. Keck, et al. were first to define HPV+ HNSCC subtypes, which they named IMS (immune strong), and CL (classical) ^15^. IMS had prominent immune and mesenchymal features while CL was enriched for putrescine (polyamine) degradation pathway. Zhang et al. re-identified the HPV+ HNSCC subtypes as IMU (immune strong) and KRT (highly keratinized) using RNA-seq and copy number variations (CNVs) ^13^, discovering a strong association between KRT and integration of HPV genes into the host genome. More recently, Locati et al. further discriminated KRT tumors into high and low stromal subtypes ^16^, and demonstrated that IMU patients have better prognosis than either high or low stromal KRT. The subtype naming convention of IMU/KRT was adopted in the Nature review, Leemans et al ^17^, and these subtypes have now been characterized using additional high-throughput technologies, including DNA methylation ^18^ showing stronger global hypomethylation in KRT, and DNA hydroxymethylation ^19^. The IMU/KRT subtypes have also been demonstrated to significantly associate with HPV E6 isoform gene expression, with KRT tending to have higher levels of the spliced E6*I isoform compared to E6 full length ^20,21^.

Although unsupervised learning methods such as clustering have the potential to reveal cancer subtypes and enable their characterization, the clusters and subtype assignments obtained naturally vary across studies. This inconsistency arises from susceptibility in the methods to factors like cohort attributes, sample quality, RNA preparation methodologies, technical variations between batches, and the specific clustering algorithm utilized ^7^. Thus, a consistent, reproducible approach is required.

To overcome the current limitations in subtyping HPV+ HNSCC tumor samples and provide a standardized subtype classification of new tumors with RNA-seq data, we trained and built a robust machine learning (ML) classifier, including several steps to enhance rigor and reproducibility. We first used 84 HNSCC HPV+ samples from two cohorts (18 from University of Michigan and 66 from TCGA) to train and test an ensemble classifier involving five ML models and three predefined gene sets as input features. We then applied our classifier to two additional cohorts of HPV+ OPSCC tumors (83 samples from the Ohio State University Comprehensive Cancer Center made available through the HPV Virome Consortium ^22^ and 62 new HPV+ OPSCC samples from the University of Michigan) and found results consistent with known subtype features and clustering results. We introduce a user-friendly webtool that streamlines and simplifies the process of subtyping HPV+ HNSCC tumors for future research. Lastly, we performed meta-analysis of the 219 subtyped HPV+ HNSCC unique patient samples and identified 21 relevant pathways and clinicodemographic variables associated with subtype.

## Methods

### HNSCC Cancer Datasets

In this study, four RNA-seq datasets were used. Two of them were 18 HPV+ HNSCC cases from University of Michigan (*UM18*) (available from GEO #GSE74956) and 66 HPV+ TCGA HNSC samples (*TCGA*), which we used to train and test the classifier, as their subtypes were previously identified (n=84: 33 IMU & 51 KRT)^13^. The other two datasets, which we utilized for validation, were from the HPV Virome Consortium (*HVC*) (n=83 HPV+; available from European Genome-phenome Archive EGAD00001004366) and a newly introduced University of Michigan (UM) OPSCC cohort (*UM67*) (n=62 HPV+) from which we used RNA from formalin-fixed, paraffin-embedded (FFPE) blocks. Written informed consents were obtained and the study was approved by the University of Michigan Institutional Review Board (See *Supplementary Methods*). Ten samples (duplicated10) in UM67 had matched FF samples in UM18. Raw gene counts of all 229 samples from the four cohorts were converted to log_2_CPM values and then normalized by each gene (z-transformed) for training, testing and validating purposes.

### Feature Selection

For training, we designed three varying-sized gene sets; the smallest one was derived from KECK (IMS/CL^15^) and the two larger sets were obtained from the Zhang et al (IMU/KRT^13^). Each training gene set was selected from the most differentially expressed genes between subtypes and balanced by IMU/KRT differentially expressed pathways (See *Supplementary Methods*). Ultimately, we obtained gene sets of size 10 (KECK), 50 (IMU_KRT_small), and 148 (IMU_KRT_large) (*Supplementary Table S1*).

Either the Z-scores of the three pre-selected gene sets (referred to as Non-PCA), or principal component analysis embeddings covering 80% of the total variance of these matrices (referred to as PCA), served as the training features.

### Classifier model training

To improve the performance and robustness of our model, we used an ensemble approach by training five different ML models and applying majority voting on the 15 (5 ML methods × 3 feature gene sets) individual models to make the final prediction. We tuned the ML models’ hyperparameters by 5-fold cross-validation (CV) (see *Supplementary Methods*). One ensemble model was trained for each format of input features (PCA and Non-PCA).

### Validation of the ensemble ML subtype classifier

To validate the robustness of the ensemble classifier, we applied it on two additional independent HPV+ OPSCC cohorts (UM67 and HVC) and checked whether 1) the classifier results for the duplicated10 samples matched the original subtype for those patients; 2) the assigned subtypes correlated well with the clinical characteristics, molecular features and pathway scores known to be associated with subtype; and 3) the classifier results were consistent with results from unsupervised clustering utilizing all genes differentially expressed by subtype (see *Supplementary Methods*).

### Webtool development and usage

A user-friendly webtool was developed accepting a matrix of gene counts or log_2_CPM values. We provide our HPV+ UM18 cohort to assist in mitigating technical batch effects between training data and user input, and to assure accurate results for small numbers of samples. See *Supplemental Methods* for more details.

### Calculation of molecular variables

For all samples, HPV+ HNSCC-relevant gene expression signature scores were calculated to characterize tumor immune microenvironment, differentiation state, HPV gene activity, and oxidation-reduction. These were generated as sample-wise pathway scores aggregated by the rank of log_2_CPM gene expression levels. First, for each gene in the relevant pathway, we ranked the samples according to their expression levels. For each sample, the ranks of the genes were summed, and the resulting values were z-score transformed across samples. To calculate the epithelial-to-mesenchymal transition (EMT) score ^23^, negatively regulated genes were also ranked in descending order. The gene sets used are in *Supplementary Table S2*.

The following scores were calculated as previously published: radio resistance score using logTPM values ^24^, which were calculated using the DGEobj.utils version 1.0.6 convertCounts function, the E6 score^25,13^, the ratio of full-length E6 to all expressed E6 isoforms (E6_FL_/E6_ALL_) influence score ^21^, and the E6_FL_ Activation score^25^. Cell type proportions were calculated using CIBERSORTx cell type deconvolution, and HPV RNA integration events were determined using SurVirus ^26^(see *Supplemental Methods*).

### Calculation of associations among subtype, molecular, and clinico-demographic variables, and generation of network graph

To generate the association network, associations between variables were calculated including cohort as a covariate using logistic regression (categorical-categorical) or ANOVA tests (see *Supplemental Methods*).

## Results

### Ensemble classifier to subtype HPV+ HNSCC bulk RNA-seq samples

Although the IMU subtype is characterized by a strong immune response and KRT by high levels of keratinization, we found that measures of immune infiltration (T cell activation scores or T cell proportion) and keratinization scores (see Methods) were inadequate to accurately subtype tumors in UM18 and TCGA cohorts (Supplementary Figure S1 A-B). This motivated us to develop a ML-based IMU/KRT subtype classifier, taking several steps to ensure its rigor and reproducibility at each phase: training, testing, and application (Supplementary Table S3).

To select training features, we used top differentially expressed genes between subtypes (*see* Methods). We designed three varying-sized gene sets (10, 50, and 148 genes) to balance the risk of overfitting with including sufficient information. (Figure 1A, Supplementary Table S1, Supplementary Figure S2A, see Methods). As a sanity check, we verified that the three gene sets were able to effectively separate TCGA samples by subtype using standard PCA, indicating their potential as training features (Supplementary Figure S2 B-D).

**Figure 1:**
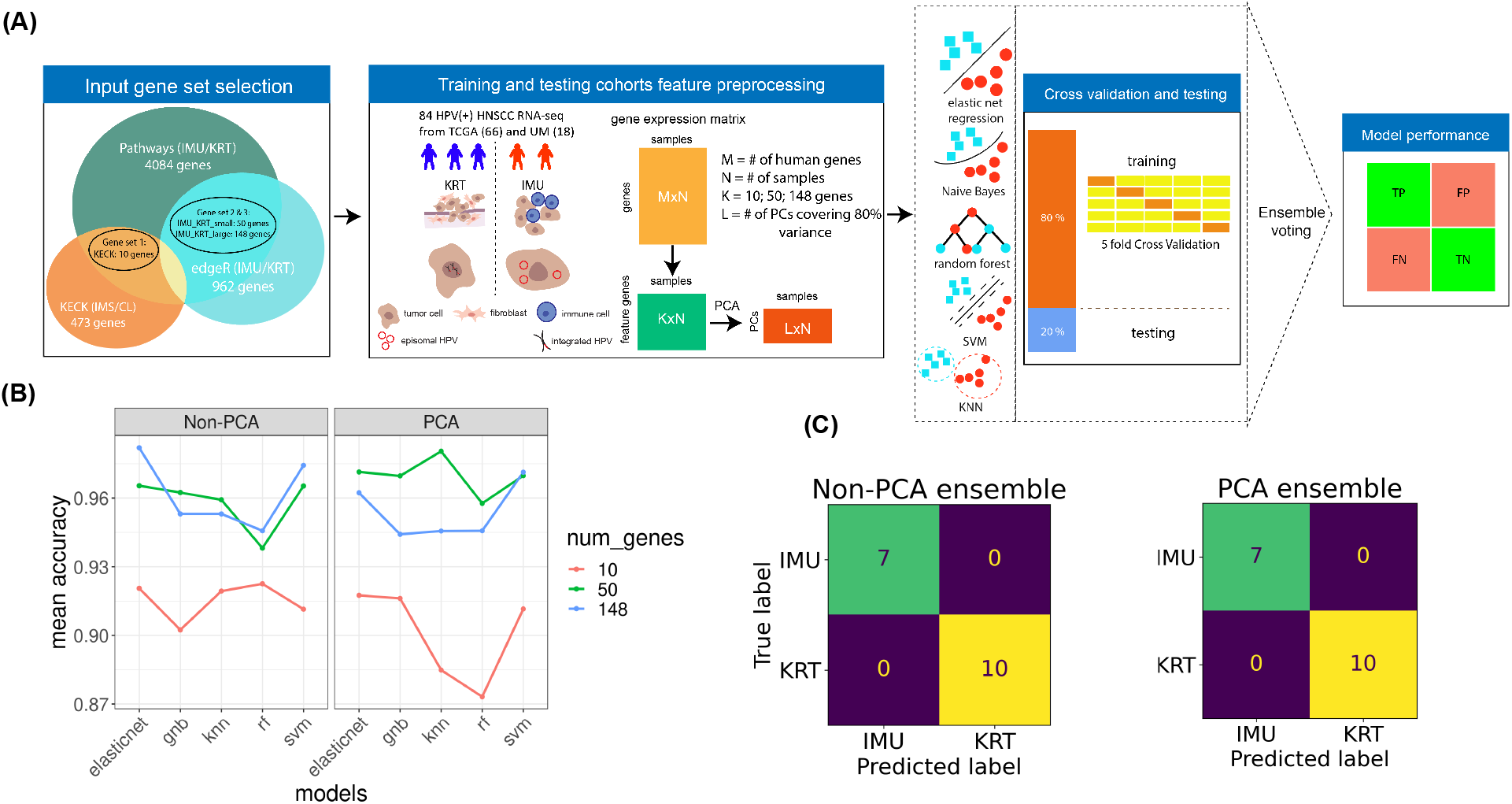
Ensemble classifier training, cross validation, and testing. (A) The schematic description of input gene set selection, pre-processing for training and testing, cross validation and testing, and implementation of ensemble model. (B) The mean cross validation (CV) accuracy for each ML model and input gene set from training. (C) Confusion matrices from test results for non-PCA and PCA based ensemble model.

After feature selection, we trained the classifier on UM18 and 49 of the TCGA cohort (n=67;18+49). To enhance the robustness of the classifier, we used an ensemble approach embedding five ML algorithms and three input gene sets in two feature formats (PCA and Non-PCA) (Figure 1A, See Methods). By comparing mean cross validation (CV) accuracy, which represents model robustness, between PCA and Non-PCA, we did not observe a significant difference (Figure 1B). We found that models using more genes (50 and 148 genes) tended to have ∼0.96 mean CV accuracy compared to ∼0.92 for the gene set of 10, indicating improvement for this dataset (Figure 1B), but this benefit may be offset by overfitting in other datasets. The fluctuations in mean CV accuracy across ML algorithms demonstrated that no single ML algorithm was overall optimal (Figure 1B, Supplementary Figure S3A-B). The observed variabilities demonstrate the value of an ensemble approach by reducing bias from a single gene set or a single ML algorithm.

We evaluated our ensemble model by testing it on the remaining TCGA samples (7 IMU; 10 KRT). Based on the confusion matrices, (Supplementary Figure S3C-D), individual ML misclassification occurred in both directions (IMU to KRT and KRT to IMU), indicating no bias toward either subtype. Although single-model misclassifications occurred for six samples across models and input gene sets (Supplementary Table S4), the final ensemble model achieved 100% accuracy for both the PCA and Non-PCA format (Figure 1C).

### Application of the subtype classifier to two additional HPV+ OPSCC cohorts validates its accuracy and robustness

To further validate the robustness and generalizability of our subtype classifier, we applied it to RNA-seq data from two additional HPV+ OPSCC cohorts (83 HVC & 62 UM67 samples) and assessed it from four perspectives. Overall, 101/145 (70%) of the samples had majority votes of 15 versus 0 or 14 vs 1, while only 11 samples had votes split by 6 vs 9 or 7 vs 8 in both PCA and Non-PCA results (Supplementary Table S4). First, we examined *duplicated10* results (See Methods) and found all ten predictions were consistent with the original UM18 pre-defined subtypes (Supplementary Table S4), confirming that the classifier is robust within patients and across source of biospecimens (FFPE versus FF).

We next examined the expression of 24 IMU/KRT differential genes, which were previously-selected ^13^ to represent the five key differential pathways between subtypes, in all newly classified samples. Importantly, only three of the 24 (SFN, HLA-DQB2, BCL2) were used in training. For both the UM67 and HVC cohorts, the Non-PCA and PCA version of the classifier subtyped the samples in close agreement with expected changes in these genes (Figure 2B, Supplementary figure S4A). The PCA-based classifier results were highly consistent with those from Non-PCA, confirming the accuracy of the classifier for both input formats. Based on these results, one can see that the classifier can identify minority cases having both IMU and KRT characteristics using the voting pattern. For instance, HVC samples GS18070 and GS18034 had inconsistent assignments between PCA and Non-PCA (Figure 2B), and their majority voting results were 6 versus 9 and 7 versus 8, respectively, with predicted IMU probabilities between 0.4 and 0.6 (Supplementary Table S4).

**Figure 2:**
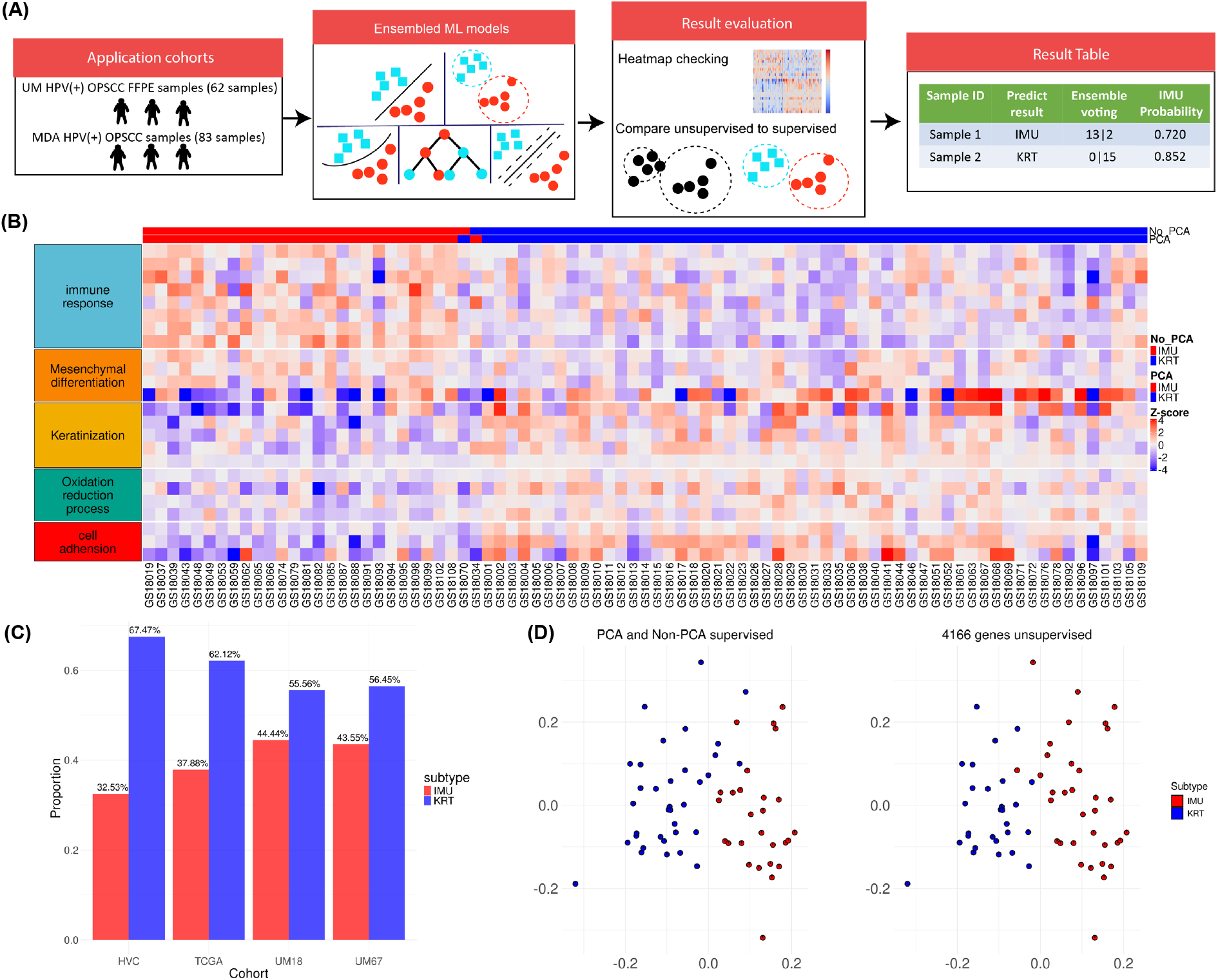
Application of the classifier on two independent HPV+ OPSCC cohorts validates the classifier accuracy and robustness. (A) The schematic for applying the classifier on two independent patient cohorts and evaluating the classifier performance. (B) Heatmap showing that 24 key genes in pathways separate the IMU/KRT subtypes for the *HVC* cohort, with PCA and Non-PCA predicted results ordered as annotation rows. (C) Proportion of the final subtypes for each cohort of patients, UM18 and UM67 have around 55% KRT while HVC and TCGA cohorts have ∼65% KRT. (D) PCA visualization of genes from union gene sets (KECK, IMU versus KRT paper and edgeR genes, see Supplementary Methods) showing differences in the separation of the subtypes for the *UM67* cohort, colored by subtype prediction results from PCA and Non-PCA classifier (top) and unsupervised (bottom).

We next validated the robustness of the classifier by testing random subsets of the features (30%, 50% and 80%), and found that 30% led to eight (5.5%) samples being misclassified, whereas 50% led to five (3.4%) being misclassified, and at 80% only two (Supplementary Figure S5 A,B). These results provide an estimate of accuracy for various levels of missing data. For all cohorts involved in this study, we consistently found around 60% of patients were KRT and 40% were IMU (Figure 2C), confirming stability of the classifier.

As a final assessment of classifier performance, we compared the Non-PCA classifier-assigned subtypes with unsupervised clustering results of all previously found DEGs by subtype (4166 genes) (See *supplementary methods*). Overall, 78% of the samples from HVC (64/83) and 88% from UM67 (55/62) had consistent results. Samples with contradictory results tended to be at the boundary in PCA visualization (Figure 2D and Supplementary Figure 4B). These results demonstrate the inconsistency of relying on unsupervised learning for subtyping tumors. To further examine the inconsistent samples, we investigated genes in the IMU/KRT differential pathways and found that unsupervised clustering tended to cluster more samples to IMU for this cohort, and samples inconsistent between unsupervised clustering and classifier results displayed pathways characteristics of both subtypes (Supplementary Figure S4C-D). The classifier tended to result in near-balanced votes or uncertain IMU probabilities for these samples (Supplementary Figure S4C-D), further demonstrating how users will be able to determine which samples do not align neatly with either IMU or KRT.

### Subtype is central to biological and clinically-relevant HPV+ cancer characteristics

To illustrate the importance of molecular subtype in HPV+ HNSCC tumor research and translational studies, we tested for significant associations among subtype and 37 carefully selected clinical, demographic, and molecular variables relevant to HPV+ HNSCC using all four cohorts. Of the variables tested, 22/37 (59.4%) were available and calculated for all cohorts. Overall, 21 variables were significantly associated with subtype (Figure 3A).

**Figure 3:**
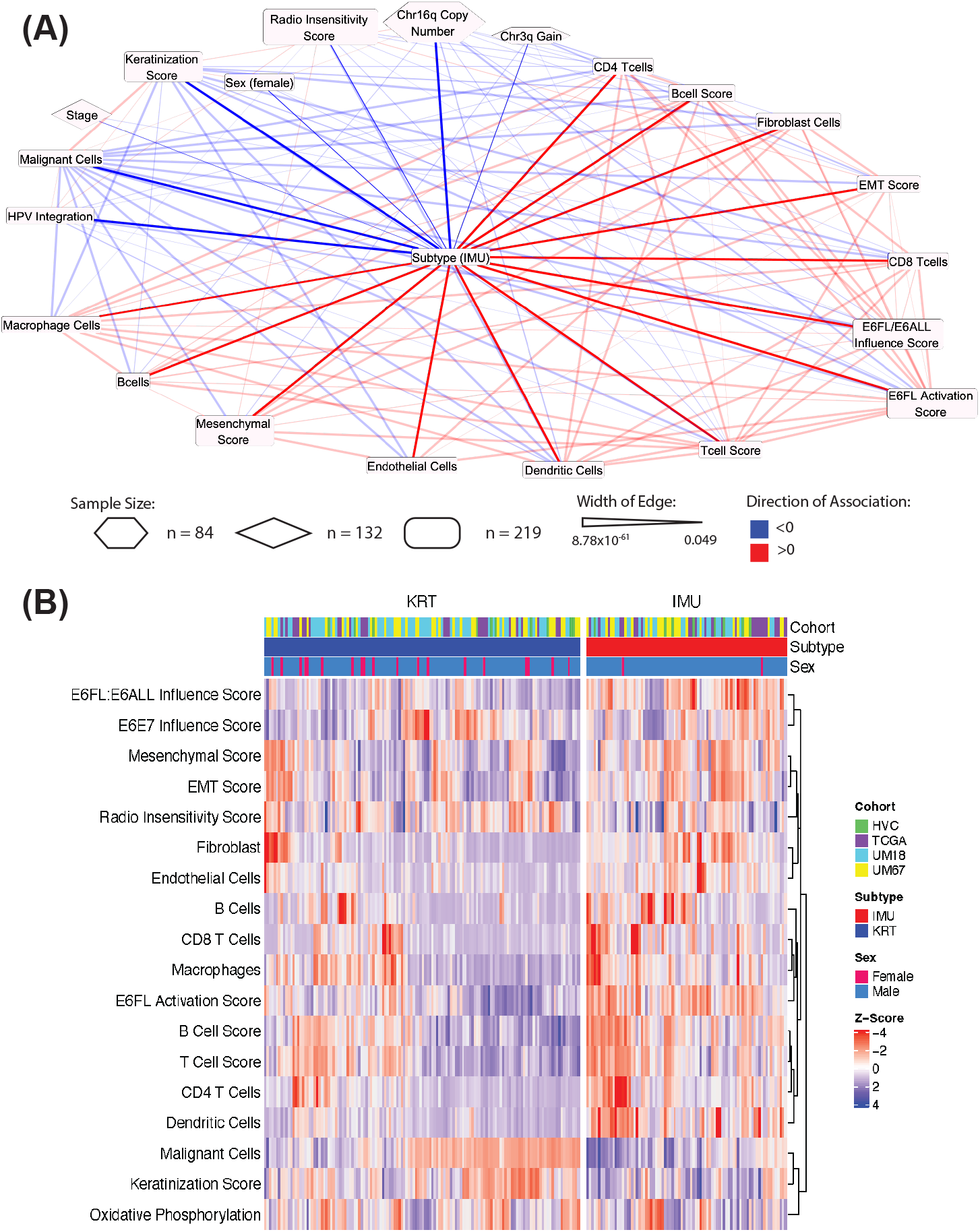
Subtype is central to biological and clinically relevant HPV+ cancer characteristics. (A) Network of clinicodemographic variables significantly associated with subtype. Red edges represent positive associations with IMU and blue edges represent positive associations with KRT. Edges between subtype and other nodes are bold for ease of viewing. The width of the edges represents the strength of association by p-value – wider being more highly significant. The shape of the nodes represents the size of the cohort used.(B) Heatmap made with R package ComplexHeatmap using z-scores of gene expression corrected for cohort effect with linear regression. Gene set scores and deconvolution derived cell-type proportions were clustered with Euclidean distance and compete linkage clustering was used on samples.

Previously known associations with subtype were reconfirmed including the association of IMU tumors with stronger EMT (p-value: 2.25×10^−4^), lower Chr16q copy number (p-value: 3.70×10^−6^), and heightened immune response as demonstrated by the significant associations with macrophage cells (p-value: 6.59×10^−4^), B cells (p-value: 1.73×10^−4^), B cell activation score (p-value: 8.67×10^−9^), dendritic cells (p-value: 1.58×10^−11^), T cell activation score (p-value: 2.18×10^−6^), CD8+ T cells (p-value: 4.62×10^−4^), and CD4+ T cells (p-value: 5.48×10^−6^) (Figure 3B). Also associated with IMU was the E6 full length (E6_FL_) activation score (p-value: 1.61×10^−12^) ^25^, which estimates the activity level of the HPV oncogene E6. The E6 full length ratio (calculated as E6_FL_/E6_ALL_) influence score ^21^ (p-value: 1.18×10^−10^) represents the influence of E6_FL_/E6_ALL_, or the proportion of total E6 that is the full length isoform as opposed to its spliced forms denoted by E6*. Thus, KRT tumors had more E6* influence. We reconfirmed the associations of KRT tumors with heightened keratinization (p-value: 2.53×10^−8^), a high probability of expressed HPV integration (p-value: 3.53×10^−6^), and copy number gains in Chr 3q (p-value: 0.011). In addition, we performed recurrence analysis with the UM67 cohort, finding that KRT patients were more likely to recur when controlling for overall stage (p=0.08027; HR=0.23) (Figure 4A). Additionally, we found that KRT tends to have a higher estimated radiation resistance (p-value: 0.0050) than IMU (Figure 4B) and higher AJCC clinical stage (p-value: 0.039) (Figure 3C). We found a novel association between sex and subtype (p-value: 0.0082) demonstrating females are more likely to be KRT than IMU (Figure 4D). No association was found between subtype and p53 mutational status, drinking status, smoking status, packs per year smoked, genomic instability, N stage, T stage, respiratory electron transport chain, 3-year survival, ACE score, or oxidative phosphorylation (Supplementary Table 5).

**Figure 4:**
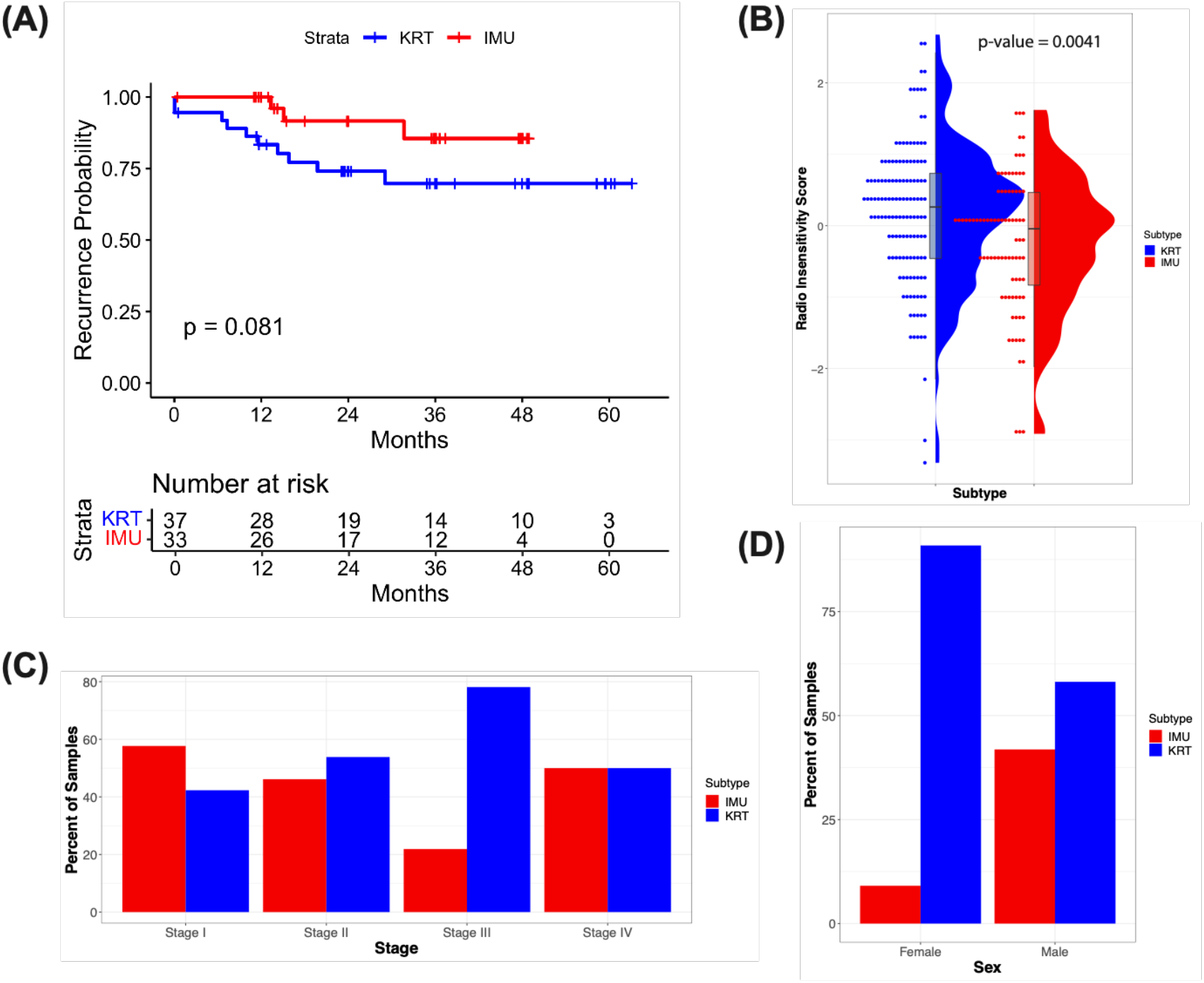
(A) Kaplan-Meier curve of recurrence probability for IMU vs. KRT. P-value was calculated by log-rank test. (B) Raincloud plot of subtype and radio insensitivity score corrected for cohort effect. P-values were calculated by t-test. (C) Bar plot of stage by subtype. (D) Bar plot of sex by subtype.

### Implementation of the subtype classifier

The classifier is available as a python-based model on GitHub (https://github.com/shengzhulst/IMUKRTclassifier), and as a webtool (https://hpv-hnscc-subtypeclassifier.dcmb.med.umich.edu/). The webtool can remove batch effects among the input samples; or between input and our UM18 cohort. The classification results are displayed as tables, PCA visualizations and an interactive heatmap to help the user evaluate the classifier’s performance (See Methods).

## Discussion

Similar HPV+ HNSCC subtypes have been rediscovered multiple times by gene expression-based unsupervised clustering and DNA methylation based deconvolution^11^. Consistently, HPV+ HNSCC tumors have been characterized as either immune strong (IMU) or highly keratinized (KRT)^13^. These subtypes have been further characterized based on mutations, CNVs, DNA methylation^18^, DNA hydroxymethylation^19^, and comprehensively reviewed in Qin et al ^11^. However, previous classifications have not provided the ability to classify new tumors. The IMU/KRT framework described here builds on existing knowledge of HPV+ HNSCC phenotypes and provides a method for classifying future samples which will aid researchers in disentangling HPV+ HNSCC tumor heterogeneity.

After classifying 219 tumors as IMU/KRT, we sought to understand features significantly correlated with each subtype. Our association tests confirmed that KRT is more likely to have chr3q gains, which is where the gene *PIK3CA* resides, and that IMU is more likely to have loss of chr16q, where several tumor suppressor cadherin genes reside including E-cadherin and P-cadherin. This is consistent with previous findings that KRT is more likely to have activating *Pik3ca* mutations and that IMU is associated with an EMT signature where a switch from E-cadherin to N-cadherin occurs. Our findings also revealed that females are more likely be classified as KRT, and KRT tumors tend to be more radiation resistant. Radiation resistance was estimated using a well-established pan-cancer gene signature^24^. Additionally, we found that KRT tumors tend to be higher stage which is supported by previous findings that KRT tumors result in worse survival. Although Locati et al found that patients with IMU-like tumors have better survival than KRT-like tumors (defined as CI1 (immune strong) vs. CI2 and CI3 (highly keratinized))^14^, we did not find a significant association with 3-year survival. Instead, we found an association between recurrence and subtype with KRT patients being more likely to recur by approximately 75% (HR=0.23). Regardless of survival and recurrence differences, tumor subtypes may point to different tendencies in likelihood for local versus regional or distal recurrence. HPV-negative HNSCC tumors are more likely to progress due to local invasion, whereas HPV+ HNSCC tumors are more likely to progress due to distant metastasis^27^. Future work testing whether IMU tumors are more likely to lead to distant metastasis and KRT tumors to local spread would be well-motivated, given the closer overall resemblance of KRT to HPV-negative oropharynx tumors and the higher EMT signature of IMU. In addition, given the lower radiation resistance signature and high immune cell infiltration of IMU tumors, one may hypothesize that IMU patients with N0 nodal status may be candidates for de-escalation trials. Unsupervised clustering applied to HPV+ HNSCC RNA-seq identified generally reliable subtypes, but it lacked reproducibility^7,11^. To overcome this, we took many steps in addition to using a supervised classifier, as outlined in Supplementary Table S3. We implemented multiple ML algorithms, used multiple input gene sets to increase generality and robustness, trained on data from two cohorts, and tested the consistency between FFPE and FF samples. To minimize batch effects, we use Combat-seq ^28^, and provide a core set of samples to offset batch effects in new samples. Finally, we assessed the classifier in new cohorts and demonstrated classifier stability.

67 samples were used for training which is admittedly small and represents on limitation of this study. However, the steps above to enhance robustness compensated for this, and we benefitted from the relative ease that the subtypes can be distinguished, as observed by the thousands of differential expressed genes and consistent rediscovery of the subtypes. For the validation cohorts (UM67, HVC), we lacked complementary assays that could support or expand our work, and clinical data was not available for HVC. Finally, ∼10∼20% of samples could not be clearly classified, which may be due to these tumors expressing a combination of IMU and KRT phenotypes, or expressing a third rarer phenotype which cannot be easily disentangled.

Using bulk RNA-seq to study cancer subtypes is effective but is limited in terms of studying within-tumor heterogeneity and its effects. Single cell and spatial transcriptomics with the combination of multi-omics and clinical data is cost prohibitive for large cohorts but will provide us new opportunities to understand tumor subtype formation, heterogeneity, and within-tumor correlates ^29,30^. For many cancer types, subtyping facilitated unraveling the carcinogenic process, for instance, sonic hedgehog (SHH)-driven medulloblastoma with mutant TP53 always displayed chromothripsis ^31^. We hope that use of this classifier will facilitate future such discoveries for HPV+ HNSCC that may one day assist in treatment decisions.

## Data sharing Statement

The UM67 samples raw count are available at GSExxxx (In progress), with aligned bam files are uploaded to European Genome-Phenome as EGAxxxxxxx (In progress).

## Acknowledgements

This work was supported by National Institutes of Health grants R01-CA250214, T32-CA140044, and P01-CA240239. We acknowledge the Advanced Genomics Cores at the University of Michigan. This study makes use of data generated by Drs. Gillison, Symer and Akagi in the HPV Virome Consortium, formerly at the Ohio State University Comprehensive Cancer Center and now at University of Texas MD Anderson Cancer Center. Funding and computational support for these data were provided by the Ohio State University Comprehensive Cancer Center, the Ohio Supercomputer Center, Cancer Prevention & Research Institute of Texas and the University of Texas MD Anderson Cancer Center.

## References

1. Meacham CE, Morrison SJ. Tumour heterogeneity and cancer cell plasticity. Nature. 2013;501(7467):328–337.

2. Yersal O, Barutca S. Biological subtypes of breast cancer: Prognostic and therapeutic implications. World J Clin Oncol. 2014;5(3):412–424.

3. Rudin CM, Poirier JT, Byers LA, et al. Molecular subtypes of small cell lung cancer: a synthesis of human and mouse model data. Nat Rev Cancer. 2019;19(5):289–297.

4. Collisson EA, Bailey P, Chang DK, Biankin AV. Molecular subtypes of pancreatic cancer. Nat Rev Gastroenterol Hepatol. 2019;16(4):207–220.

5. Marisa L, de Reyniès A, Duval A, et al. Gene expression classification of colon cancer into molecular subtypes: characterization, validation, and prognostic value. PLoS Med. 2013;10(5):e1001453.

6. Zhao L, Lee VHF, Ng MK, Yan H, Bijlsma MF. Molecular subtyping of cancer: current status and moving toward clinical applications. Brief Bioinform. 2019;20(2):572–584.

7. Källberg D, Vidman L, Rydén P. Comparison of methods for feature selection in clustering of high-dimensional RNA-sequencing data to identify cancer subtypes. Front Genet. 2021;12:632620.

8. Almarzooqi S, Hashim MJ, Awwad A, Sharma C, Saraswathiamma D, Albawardi A. Lower prevalence of human Papillomavirus in Head and neck squamous cell carcinoma in middle eastern population: Clinical implications for diagnosis and prevention. Cureus. 2023;15(2):e34912.

9. Menezes FDS, Fernandes GA, Antunes JLF, Villa LL, Toporcov TN. Global incidence trends in head and neck cancer for HPV-related and -unrelated subsites: A systematic review of population-based studies. Oral Oncol. 2021;115(105177):105177.

10. Lechner M, Liu J, Masterson L, Fenton TR. HPV-associated oropharyngeal cancer: epidemiology, molecular biology and clinical management. Nat Rev Clin Oncol. 2022;19(5):306–327.

11. Qin T, Li S, Henry LE, Liu S, Sartor MA. Molecular tumor subtypes of HPV-positive head and neck cancers: Biological characteristics and implications for clinical outcomes. Cancers (Basel). 2021;13(11):2721.

12. Lewis JS Jr, Mirabello L, Liu P, et al. Oropharyngeal squamous cell carcinoma morphology and subtypes by human Papillomavirus type and by 16 lineages and sublineages. Head Neck Pathol. 2021;15(4):1089–1098.

13. Zhang Y, Koneva LA, Virani S, et al. Subtypes of HPV-positive head and neck cancers are associated with HPV characteristics, copy number alterations, PIK3CA mutation, and pathway signatures. Clin Cancer Res. 2016;22(18):4735–4745.

14. Locati Serafini, Iannò, et al. Mining of self-organizing map gene-expression portraits reveals prognostic stratification of HPV-positive head and neck squamous cell carcinoma. Cancers (Basel). 2019;11(8):1057.

15. Keck MK, Zuo Z, Khattri A, et al. Integrative analysis of head and neck cancer identifies two biologically distinct HPV and three non-HPV subtypes. Clin Cancer Res. 2015;21(4):870–881.

16. Sahovaler A, Kim MH, Mendez A, et al. Survival outcomes in human Papillomavirus-associated nonoropharyngeal squamous cell carcinomas: A systematic review and meta-analysis. JAMA Otolaryngol Head Neck Surg. 2020;146(12):1158–1166.

17. Leemans CR, Snijders PJF, Brakenhoff RH. The molecular landscape of head and neck cancer. Nat Rev Cancer. 2018;18(5):269–282.

18. Qin T, Li S, Henry LE, et al. Whole genome CpG-resolution DNA methylation profiling of HNSCC reveals distinct mechanisms of carcinogenesis for fine-scale HPV+ cancer subtypes. Cancer Res Commun. Published online August 7, 2023. doi:10.1158/2767-9764.crc-23-0009

19. Liu S, de Medeiros MC, Fernandez EM, et al. 5-Hydroxymethylation highlights the heterogeneity in keratinization and cell junctions in head and neck cancers. Clin Epigenetics. 2020;12(1):175.

20. Koneva LA, Zhang Y, Virani S, et al. HPV integration in HNSCC correlates with survival outcomes, immune response signatures, and candidate drivers. Mol Cancer Res. 2018;16(1):90–102.

21. Qin T, Koneva LA, Liu Y, et al. Significant association between host transcriptome-derived HPV oncogene E6* influence score and carcinogenic pathways, tumor size, and survival in head and neck cancer. Head Neck. 2020;42(9):2375–2389.

22. Symer DE, Akagi K, Geiger HM, et al. Diverse tumorigenic consequences of human papillomavirus integration in primary oropharyngeal cancers. Genome Res. 2022;32(1):55–70.

23. Zeisberg M, Neilson EG. Biomarkers for epithelial-mesenchymal transitions. J Clin Invest. 2009;119(6):1429–1437.

24. Dai YH, Wang YF, Shen PC, et al. Radiosensitivity index emerges as a potential biomarker for combined radiotherapy and immunotherapy. NPJ Genom Med. 2021;6(1):1–10.

25. Duffy CL, Phillips SL, Klingelhutz AJ. Microarray analysis identifies differentiation-associated genes regulated by human papillomavirus type 16 E6. Virology. 2003;314(1):196–205.

26. Rajaby R, Zhou Y, Meng Y, et al. SurVirus: a repeat-aware virus integration caller. Nucleic Acids Res. 2021;49(6):e33–e33.

27. Sacks R, Law JY, Zhu H, et al. Unique patterns of distant metastases in HPV-positive head and neck cancer. Oncology. 2020;98(3):179–185.

28. Zhang Y, Parmigiani G, Johnson WE. ComBat-seq: batch effect adjustment for RNA-seq count data. NAR Genom Bioinform. 2020;2(3):lqaa078.

29. Lawson DA, Kessenbrock K, Davis RT, Pervolarakis N, Werb Z. Tumour heterogeneity and metastasis at single-cell resolution. Nat Cell Biol. 2018;20(12):1349–1360.

30. Ayton SG, Pavlicova M, Robles-Espinoza CD, Tamez Peña JG, Treviño V. Multiomics subtyping for clinically prognostic cancer subtypes and personalized therapy: A systematic review and meta-analysis. Genet Med. 2022;24(1):15–25.

31. Schneider G, Schmidt-Supprian M, Rad R, Saur D. Tissue-specific tumorigenesis: context matters. Nat Rev Cancer. 2017;17(4):239–253.

