## Supplementary materials for "Tumor Subtype Classification Tool for HPV-associated Head and Neck Cancers"

**Supplementary methods:**

*Collection, RNA sequencing, and preprocessing of UM67 OPSCC University of Michigan tumors*

FFPE blocks were collected from tumors originating in the oropharynx of patients who were part of the NCI-funded UM Head and Neck (H&N) SPORE population. These patients were enrolled in the UM H&N SPORE between 2008 – 2014 and have extensive epidemiologic data and follow-up information for survival and local, regional, and distant recurrences. H&E slides were sectioned from each FFPE block and assessed by a board certified, HNC expert pathologist for degrees of cellularity and necrosis. Tissue was scraped from areas of >70% cellularity from 5-micron sections for RNA isolation. After RNA extraction via the RNAStorm FFPE Kit (Cell Data Sciences) and quality assessment via the Agilent Bioanalyzer, the QIAGEN FastSelect kit and New England Biolabs (NEB) Next Ultra II Directional RNA Library Prep Kit were used for library prep.

Four batches of RNA samples (2536-KZ, 3687-SS, 3860-SS, 4165-SS) were sequenced from 2021 to 2022. Bulk RNA-seq was generated at the UM Advanced Genomics Core with NovaSeq S4 (300 cycle and 150bp paired-end). For different batches, different amounts of RNA and cycles of PCR were performed. (2536-KZ: 61ngx14cycle, 3687-SS: 45ngx14cycle, 3860-SS: 45ngx14cycle and 4165-SS: 230x12 cycle).

*Data preprocessing of UM67 and HVC HPV+ OPSCC samples*

For data processing, Quality control was performed by FastQC 0.11.9, and adapter sequences were trimmed using Cutadapt 3.4 ^1^. The high-quality reads were then aligned to the self- built reference genome (hg38 plus high risk HPVs (HPV16, HPV18, HPV31, HPV33, HPV35, HPV39, HPV45, HPV51, HPV52, HPV56, HPV58, HPV59, HPV66, HPV68, HPV73, HPV82)) using STAR 2.7.9a ^2^, and expression levels of human genes were quantified using htseq-count 0.13.5 ^3^. Only samples with more than 200 HPV reads were regarded as HPV+ HNSCC samples.

The BAM files obtained from EGAD00001004366 (HCV cohort) were converted back to fastq with samtools v1.12 for preprocessing. Removing samples not defined as HPV+ by RNA expression resulted in 83 HPV+ HVC samples and 62 HPV+ UM67 samples. Ten HPV+ samples from nine unique patients in the UM67 cohort (FFPE) were from patients also in UM18 (FF) (named as duplicated10) (Supplementary Table S3), allowing us to directly assess the reproducibility and concordance of the subtype classifier between fresh frozen and FFPE-derived tumor samples. Staging for all samples was done using AJCC tumor stage V8 ^4^.

Raw human gene counts of all 229 RNA-seq samples were converted to log2CPM values utilizing the edgeR v3.34.1 cpm function and then normalized for each gene (z-transformed) by the scikit-learn v1.2.2 StandardScaler function with default parameters (z-score) for training, testing and validating purposes. We also implemented Combat-seq to remove batch effects from within UM67 (FFPE) and between the UM18 (FF) and UM67 cohorts.

*Preparation of input gene sets for classifier training*

To avoid data leakage, no information from any test set samples was used to define the input training gene sets. The raw gene counts from UM18 (8 IMU+10 KRT) were obtained from the IMU/KRT subtype defining paper ^5^ and used to perform differential gene expression analysis between subtypes by edgeR 3.34.1. Age was included as a covariate, and glmQLFit and glmQLFTest were applied. Genes with FDR<0.05 and absolute value of log fold change > 2 were recognized as significantly differentially expressed (962 genes, referred to as edgeR genes).

To enhance the generalizability of our study, we also included genes from another previous HNSCC subtype study based on microarray data (KECK) ^6^. We downloaded their defined subtypes (IMS/CL/BA) centroid feature genes (821 genes), and kept genes that had absolute values of difference between HPV+ related IMS and CL subtypes centroid larger than 0.4 (KECK genes: 473 genes) as differentially expressed genes for downstream analysis.

The largest gene set, which we used for assessment in this paper, was also from Zhang et al. 2016 (pathway genes: 4084 genes) ^5^. It includes all genes in IMU versus KRT differentially expressed pathways in four groups: KRT-down: Extracellular matrix, Mesenchymal differentiation, Angiogenesis; IMU-down: Epidermal cell differentiation, Oxidative reduction process, Ribosome biogenesis; KRT-up: Keratinization, Oxidation reduction process, DNA repair; IMU-up: Adaptive immune response, Innate immune response, T and B cell activation.

The union of all three gene sets (edgeR, KECK, and pathway genes (Union: 4166 genes), which represents the most comprehensive set of subtype-related genes, was used to implement unsupervised clustering tasks, and results from unsupervised clustering were further used to evaluate the subtype classification results.

We designed three training gene sets of 10, 50, and 150 genes, with the smallest set derived from KECK and the other two from edgeR. Genes in each set are from the overlapping genes between KECK or edgeR and pathway genes prioritized by their rank in terms of the degree of subtype difference and constrained by their assigned pathway. The degree of subtype difference was defined as the absolute value of the difference between IMS and CL subtype centroids (for KECK genes) or the absolute value of the log2 fold change (for edgeR genes). For each training gene set, we considered the balance between up- and down- regulated (IMU as reference), as well as including genes in pathways in each direction (IMU-up, KRT-down, KRT-up, IMU-down). IMU-up pathways (Adaptive immune response, Innate immune response, T and B cell activation) contain nearly twice as many genes as the other pathways (KRT-down, KRT-up, IMU-down) ^5^. Therefore, for each training gene set, we selected 40% of genes from the IMU-up pathways, and 20% of genes from each of the remaining three pathways (60% in total). The steps above guaranteed our training gene sets covered most significantly differentially expressed genes between subtypes, while achieving balance at the pathway level.

We used the scikit-learn “sklearn.decomposition.PCA” to perform dimension reduction on all training gene sets with 80% of the variance preserved.

We used 24 genes to visualize heatmaps in the main figure, 24 genes were previously mentioned ^7^ to represent five key differential pathways between subtypes (immune response, mesenchymal differentiation, keratinization, oxidation reduction, and cell adhesion).

*Unsupervised clustering of UM67 and HVC HPV+ OPSCC samples*

We employed consensus clustering on the z-scores from UM67 and HVC HPV+ OPSCC samples with the Union gene list (4166 genes) using the R package consensusclusterplus 1.56.0. We specified clusterAlg as “hc” and innerLinkage as “ward.D2” for the ConsensusClusterPlus function. The final unsupervised clustering results are from k=2.

*Hyperparameter tuning and training for ML algorithms*

To avoid data leakage, we included all UM18 samples in the training cohort and randomly split the 66 TCGA samples into 75% (n=49) for training and 25% (n=17) for testing. To avoid potentially imbalanced data due to small sample size and to maximize generalizability in the results, we carefully examined the labels of the training and test samples to ensure that they were balanced by subtype.

Each ML training process was implemented by scikit-learn v1.2.2 and calibrated by CalibratedclassifierCV. To tune the training hyperparameters for each ML algorithm used in this study, we applied the “GridSearchCV” function to perform a 5-fold cross validation and calculated the mean accuracy as the evaluation score. We focused on optimizing the n_estimators (100, 200, 500, 1000) for random forest, the n_neighbors (1 to 21) for KNN, whether to use linear SVM (linear, poly, rbf), the var_smoothing (0 to 1) for Gaussian naïve Bayes and the L1 ratio (0.2, 0.5, 0.8) for elastic net. Overall, we trained 30 ML models, using the 3 different input gene lists directly (Non-PCA) and using their PCA principal components (5 models x 3 gene lists x 2 types of features), and we performed majority voting (15 models) to decide the final predicted subtype.

*Webtool development and input data preprocessing*

The website was constructed using Django 4.1. It is designed to accept either gene count or log2cpm as input, and the transformation of those two types of values is accomplished using edgeR. The user input gene information column supports Entrez gene ID, Gene Symbol, or Ensemble Gene ID. To ensure consistency, the input gene information is standardized to the same version of Gene Symbol using HGNChelper 0.8.1 and org.Hs.eg.db.

Batch effects are removed with the ComBat-seq function in the sva R package (version 3.40.0). PCA-based classification is performed when the user provides less than 80% of the training genes, and non-PCA classification is performed otherwise. To address missing values in the user-provided gene lists, we use sample-wise KNN imputation with k=5 and weights=distance utilized KNNImputer in scikit-learn. The UM18 cohort is used to impute values of genes completely missing from the user-provided data. An interactive heatmap (generated using clustergrammer2) and PCA plots using each of the three gene lists and colored by final subtype predictions (seaborn 0.9.0), are generated to facilitate inspection of results, and a table containing subtype, ensemble vote, and subtype probability are provided to the user for download.

*Cell Type Deconvolution*

Cell type deconvolution was conducted utilizing CIBERSORTx with the single-cell RNA-seq HNSCC reference matrix (Newman et al. 2019). To address the distinct origins of the signature matrix and the mixture matrix, derived from single-cell RNAseq and bulk RNA-seq platforms, respectively, B-mode batch correction was activated to mitigate any technical variations. Quantile normalization and absolute mode were both disabled by default. The number of permutations was set to 100, meeting the recommended minimum for obtaining a robust deconvolution p-value. The resulting cell type deconvolution matrix included proportions of the following cell types for all samples: CD8+ T cells, CD4+ T cells, Fibroblasts, Macrophages, B cells, Malignant Cells, Mast Cells, Dendritic Cells, Myocytes, and Endothelial Cells.

*HPV Integration detection*

HPV RNA integration events were determined using SurVirus with default settings, utilizing the RNA-seq trimmed fastqs from all cohorts (Rajaby et al. 2021). Participants with at least one detected integration event were defined as integration-positive.

*Calculation of associations among subtype, molecular, and clinico-demographic variables, and generation of network graph*

For categorical-categorial associations, logistic regression was used considering the cohort (TCGA, UM18, HVC, or UM67) as a covariate. For all other associations, an ANOVA test was used with cohort as a covariate. Only variables significantly (p < 0.05) associated with subtype were included. Cytoscape (version 3.9.1) was used to generate the network graph.

*Recurrence Analysis*

To evaluate the recurrence probabilities over time for IMU and KRT, we employed Kaplan-Meier analysis. Kaplan-Meier recurrence curves were generated using the survival package in R (version 4.1.1). Recurrence curves were plotted for IMU vs. KRT to visualize differences in recurrence probabilities over time. The log-rank test was used to compare the recurrence curves between groups. To identify factors associated with recurrence, we utilized the Cox proportional hazards regression model. The model was specified using subtype (IMU vs. KRT) and the potential confounding variable, tumor stage.

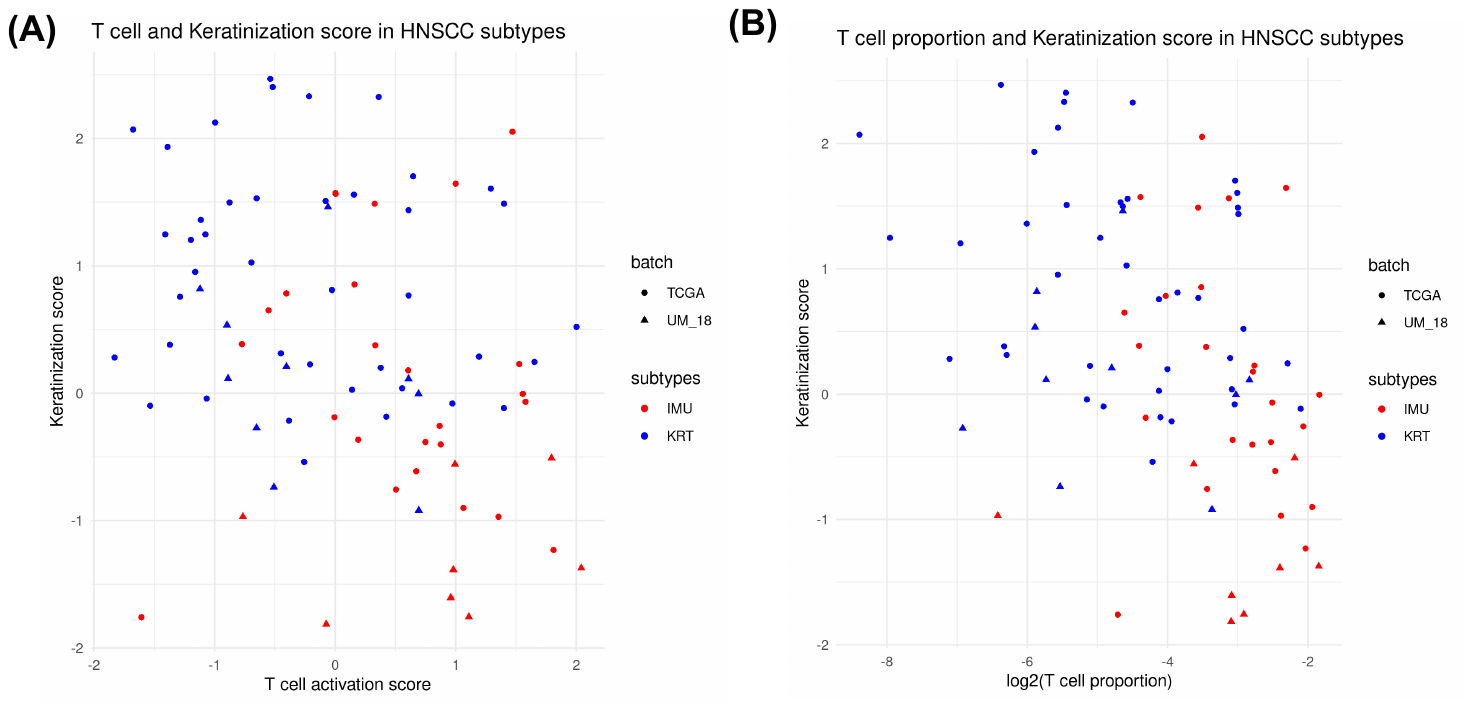

**Supplementary Figure S1:** Keratinization and T cell scores (or cell type proportion) are not sufficient to accurately classify tumors as IMU versus KRT. (A) T cell activation score versus keratinization score for the TCGA (n=66) and UM18 (n=18) cohorts (see Methods for details). (B) T cell (CD4 + CD8) proportions obtained from cell type deconvolution versus keratinization score for the same cohorts.

**Supplementary Figure S2**: Gene sets used to train the machine learning models have potential to classify subtypes. (A) Venn diagram for the three gene sets used to train the ML models (KECK: genes from Keck.et al, IMUKRT_small: smaller gene set from edgeR generated IMU versus KRT differential expressed genes, IMUKRT_large: larger gene set from IMU versus KRT differential expressed genes). (See Methods for more details). (B-D) PCA visualization for TCGA+ UM18 samples, colored by predicted subtype, using (B) KECK gene set (10 genes), (C) IMUKRT_small gene set (50 genes), and (D) IMUKRT_large gene set (148 genes).

**Supplementary Figure S3:** Confusion matrices and cross validation results for the individual ML models. (A) Non-PCA based cross validation results for each ML model and each input gene set. (B) PCA based cross validation result for each ML model and each gene set. (C) Non-PCA based confusion matrices for the testing cohort. Shown are the results for each ML model and each input gene set separately. (D) PCA based confusion matrix for each ML model and gene set combination.

**Supplementary Figure S4**: Extra evidence to support the accuracy of the classifier. (A) Heatmap shows the key genes in key pathways separate the IMU/KRT subtype for the UM67 cohort, with PCA or Non-PCA predicted results as top annotations. (B) PCA visualization of genes (4166 genes) from union gene sets (KECK, IMU versus KRT paper and edgeR based differentially expressed genes, see supplementary methods) showing differences in the separation of the subtypes for the *UM67* cohort, colored by subtype prediction results from PCA and Non-PCA classifier (top) and unsupervised (bottom). (C-D) Z-score heatmap visualization of pathway genes (See Supplementary methods) for HVC (C) and UM67 cohorts (D); Top annotation bar shows the results from unsupervised clustering based on all genes in this heatmap and supervised clustering with Non-PCA based classifier, calculated IMU subtype probability, and voting results.

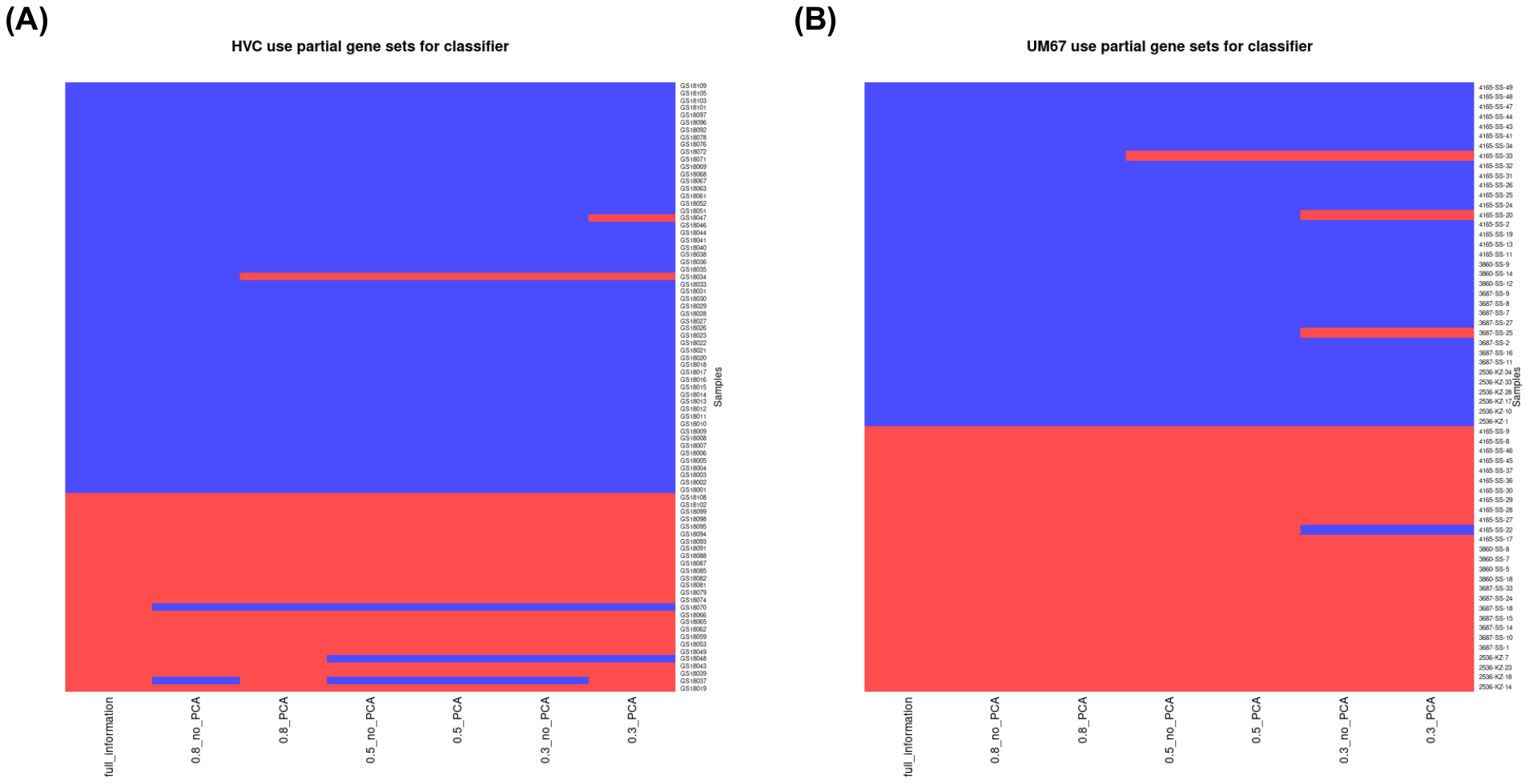

**Supplementary Figure S5**: Extra evidence to support the robustness of the classifier. (A-B) Heatmap showing the high level of classifier prediction consistency when we randomly selected 80%, 50% or 30% of genes in each training gene set for the HVC (A) and UM67 (B) cohorts.

**References**

1.	Martin M. Cutadapt removes adapter sequences from high-throughput sequencing reads. *EMBnet J*. 2011;17(1):10.

2.	Dobin A, Davis CA, Schlesinger F, et al. STAR: ultrafast universal RNA-seq aligner. *Bioinformatics*. 2013;29(1):15-21.

3.	Anders S, Pyl PT, Huber W. HTSeq—a Python framework to work with high-throughput sequencing data. *Bioinformatics*. 2015;31(2):166-169.

4.	Zanoni DK, Patel SG, Shah JP. Changes in the 8th edition of the American joint committee on cancer (AJCC) staging of head and neck cancer: Rationale and implications. *Curr Oncol Rep*. 2019;21(6):52.

5.	Zhang Y, Koneva LA, Virani S, et al. Subtypes of HPV-positive head and neck cancers are associated with HPV characteristics, copy number alterations, PIK3CA mutation, and pathway signatures. *Clin Cancer Res*. 2016;22(18):4735-4745.

6.	Keck MK, Zuo Z, Khattri A, et al. Integrative analysis of head and neck cancer identifies two biologically distinct HPV and three non-HPV subtypes. *Clin Cancer Res*. 2015;21(4):870-881.

7.	Zhang Y, Koneva LA, Virani S, et al. Subtypes of HPV-positive head and neck cancers are associated with HPV characteristics, copy number alterations, PIK3CA mutation, and pathway signatures. *Clin Cancer Res*. 2016;22(18):4735-4745.
